# Leveraging machine learning to streamline the development of liposomal drug delivery systems

**DOI:** 10.1101/2024.07.01.600773

**Authors:** Remo Eugster, Markus Orsi, Giorgio Buttitta, Nicola Serafini, Mattia Tiboni, Luca Casettari, Jean-Louis Reymond, Simone Aleandri, Paola Luciani

**Affiliations:** Department of Chemistry, Biochemistry and Pharmaceutical Sciences, University of Bern, Bern, Switzerland; Department of Chemistry and Technologies of Drugs, Sapienza University of Rome, Rome, Lazio, Italy; Department of Biomolecular Sciences, University of Urbino Carlo Bo, Urbino, PU, Italy

## Abstract

Drug delivery systems efficiently and safely administer therapeutic agents to specific body sites. Liposomes, spherical vesicles made of phospholipid bilayers, have become a powerful tool in this field, especially with the rise of microfluidic manufacturing during the COVID-19 pandemic. Despite its efficiency, microfluidic liposomal production poses challenges, often requiring laborious, optimization on a case-by-case basis. This is due to a lack of comprehensive understanding and robust methodologies, compounded by limited data on microfluidic production with varying lipids. Artificial intelligence offers promise in predicting lipid behaviour during microfluidic production, with the still unexploited potential of streamlining development. Herein we employ machine learning to predict critical quality attributes and process parameters for microfluidic-based liposome production. Validated models predict liposome formation, size, and production parameters, significantly advancing our understanding of lipid behaviour. Extensive model analysis enhanced interpretability and investigated underlying mechanisms, supporting the transition to microfluidic production. Unlocking the potential of machine learning in drug development can accelerate pharmaceutical innovation, making drug delivery systems more adaptable and accessible.

## Introduction

Liposomes, have revolutionized the field of drug delivery (Fig. 1a).^1^ Due to their versatile applications and the pandemic-driven push to standardize lipid-based delivery methods, these vesicles have garnered increasing attention.^2^ Over the last three decades, more than 14 liposome-based drug products have been approved, with applications ranging from cancer treatments to vaccines.^2^ Liposomal carriers can encapsulate both hydrophilic and hydrophobic drugs, offering a protective environment and enhancing their solubility, stability, and bioavailability.^1,3^ Further, the liposomal surface can be engineered to circulate longer in the bloodstream, improving the pharmacokinetics and biodistribution.^4,5^

**Fig. 1:**
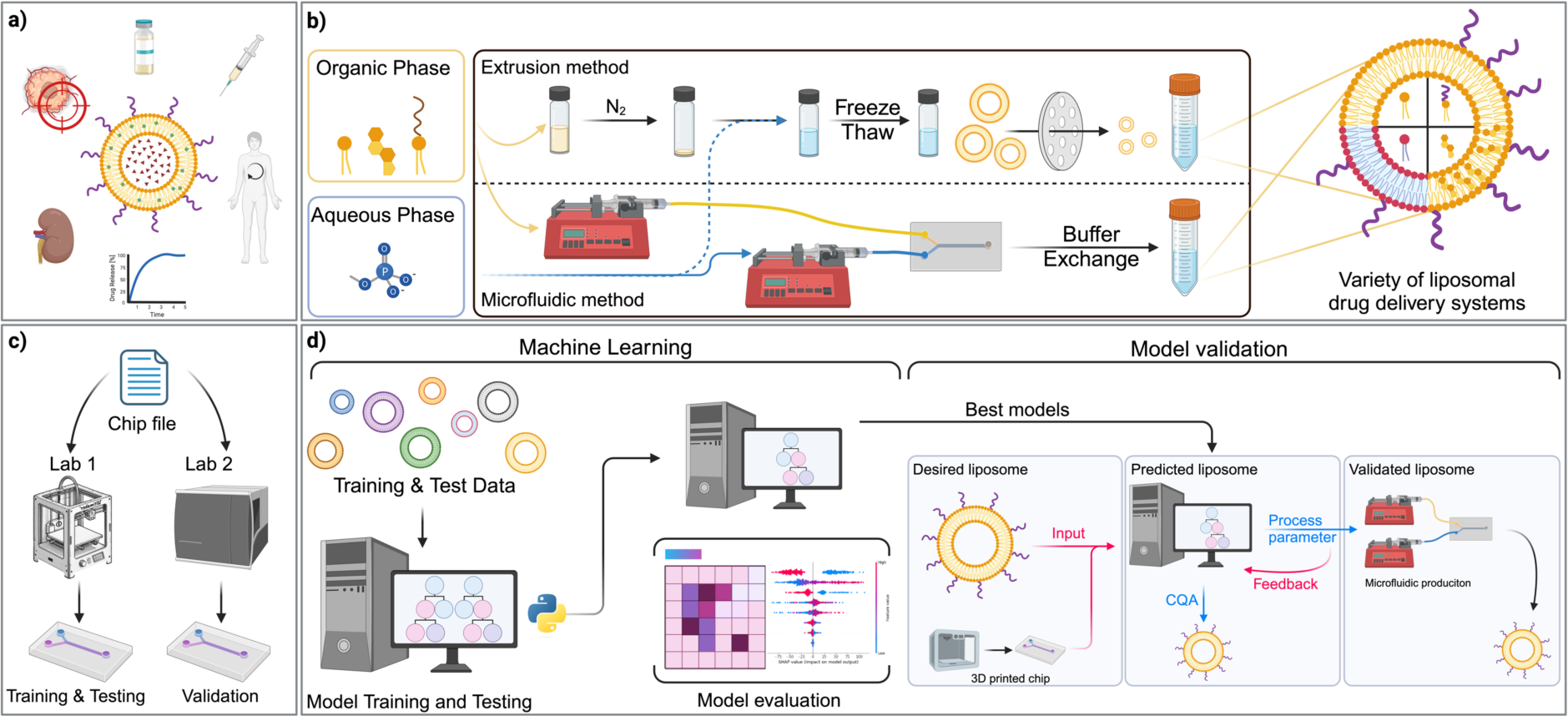
Schematic demonstration of traditional and data-driven formulation development for liposomal drug delivery systems. a) Liposomes serve as effective drug delivery systems, enhancing drug stability, enabling parenteral administration for systemic effects, and offering controlled drug release. PEGylation and precise size adjustments help reducing clearance and increase the retention of drug carriers in the body. Additionally, liposome size influences physiological processes such as hepatic uptake and tissue diffusion. b) Traditional extrusion-based versus microfluidic production method of liposomes. c) 3D printing of microfluidic chips with specific channel geometries. d) Machine learning (ML) model development, testing model selection based on model interpretation and validation in the wet lab.

Systemic drug delivery is profoundly impacted by liposome size, affecting physiological processes such as hepatic uptake, tissue diffusion, extravasation, and renal clearance.^6–8^ The size range of 50-200 nm is considered optimal for drug nanocarriers in systemic parenteral administration, balancing tissue and capillary pore size limitations.^6,9,10^ Additionally, smaller liposomes (<150 nm) demonstrate enhanced lymphatic uptake and transport, crucial for effective drug delivery.^9,11^ Further, larger liposomes might escape clearance and act as long-acting depot systems.^12,13^ Liposomes mainly consist of phosphatidylcholine (PC) lipids, essential for drug delivery due to their biocompatibility and ability to form stable bilayers.^14^ Traditional methods of liposome production, such as thin-film hydration and ethanol injection, present challenges in scalability, reproducibility, and control over liposome size and polydispersity (Fig. 1b).^15,16^ In contrast, microfluidic technology offers a promising alternative by enabling precise control over the production process. With the increasing accessibility of microfluidic devices for lipid nanoparticle (LNP) production, this technology is becoming a reality for small and medium enterprises (SMEs), encouraging formulation scientists to explore more liposome-based drug delivery systems.^15–19^ This technology involves manipulating fluids within microchannels, allowing for controlled mixing and assembly of liposomal components (Fig. 1b). Microfluidic production enhances reproducibility and scalability, generating liposomes with uniform size distributions and tailored properties, fundamental for consistent therapeutic performance.^6,15,17,20^ Additionally, it avoids chlorinating agents and special waste disposal, making it more sustainable. Further, microfluidic production offers fine control over critical process parameters (CPPs) such as flow rates, pressure, and temperature, which are crucial for optimizing liposome size, lamellarity, encapsulation efficiency, and stability.^21–23^ Despite these benefits, however, several challenges hinder clinical and industrial applications. The primary obstacle is translating bench-scale liposomal production to larger scales, as traditional methods often do not scale up seamlessly.^23^ However, transitioning to more scalable microfluidic production is not trivial as different liposomal formulations respond uniquely to microfluidic conditions, requiring extensive optimization on a case-by-case basis.^23^ Variations in microfluidic chip design further complicate standardization. Advances in 3D printing could help by enabling robust, standardized chip fabrication (Fig. 1c).^24,25^ While extrusion can control liposome size, achieving this precision in microfluidics is more complex due to the interplay of various factors, often requiring extensive experimentation to adjust the flow rate ratio (FRR; aqueous: organic flow rate) at a sufficient total flow rate.^20,23,26^

Mathematical models like Design of Experiments (DoE) have guided the development of microfluidic liposome production processes, elucidating the relationships between process parameters and the resulting liposome characteristics.^22,24,27^ However, their applicability is often limited to low-dimensional design spaces, and their scalability and generalization are restricted. In recent years, the application of machine learning (ML) and artificial intelligence (AI) in pharmaceutical sciences has gained traction due to their ability to model high-dimensional problems.^21,22,28–31^ ML is valuable throughout the entire drug discovery and development pipeline,^21,32^ from exploring chemical space to identify new hit compounds^32–36^, to predicting synthetic pathways,^37–39^ Absorption-Distribution-Metabolism-Excretion Toxicity (ADMET) properties^40,41^ and target activity.^42^ However, many methods rely on external datasets often lacking quality.^43,44^ Previous work using in-house data has demonstrated that ML can accurately learn and predict experimental outcomes with clean data, even with smaller datasets.^45^ The primary limitation for most applications so far is the scarcity of comprehensive and high-quality data. However, this does not apply to microfluidic liposome production, as a fair amount of in-house data can readily be produced. Therefore, ML holds significant promise for predicting lipid behaviour during microfluidic production with high accuracy.^22,30,31,46,47^ Despite their ability to model complex high-dimensional data, though, ML models often lack explainability, making them less suitable for gaining deeper insights.^48^ Explainable AI (XAI) techniques, allow researchers to extract information from black-box models and provide more transparency to the predictions.^48^ Combined, ML and XAI can potentially reduce reliance on empirical methods and provide increased transparency, accelerating the optimization of the liposomal production process.^30,31,46,47,49^

Herein we report on training ML models, with an in-house generated data set to accurately predict critical quality attributes (CQAs), including the formation and size of liposomes, as well as process parameters for their microfluidic production. We applied the principles of XAI, thereby, one can gain deeper insights into the complex interactions between formulation- and process-parameters, improving the prediction of lipid behaviour during microfluidic production. By training these models, we aim to establish a reliable framework that enables transitioning from traditional liposome production methods to microfluidic techniques (Fig. 1d). This capability can streamline the development of liposomal drug delivery systems, reducing reliance on empirical methods and accelerating the optimization process.^30,31,46,47^ Our ultimate goal is to develop a robust, scalable, and standardized process widely adoptable in the pharmaceutical sciences, to accelerate the delivery of therapeutic agents to clinics and patients.

## Results and Discussion

### Liposomal formulation screening

A comprehensive range of lipid-based formulations has been screened following multiple full factorial DoEs and using a microfluidic process, varying CPPs to optimize liposome characteristics. The primary CQA obtained in response to these variations was liposome size, a key determinant of their therapeutic efficacy and stability.^6^ The full factorial design allowed us to screen a vast portion of the experimental space, distributing the experimental points among the investigated CPPS’ s range. The lipid excipients selected for this study include dimyristoylphosphatidylcholine (DMPC), dioleoylphosphatidylcholine (DOPC), hydrogenated soy phosphatidylcholine (HSPC), dipalmitoylphosphatidylcholine (DPPC), and palmitoyloleoylphosphatidylcholine (POPC). This selection covers a broad spectrum of lipids commonly used in the development of liposomal drug delivery systems, providing a diverse basis for the analysis.^14^ The chosen lipids are pivotal for subcutaneous (s.c.) and parenteral drug delivery, given their distinct biophysical properties and interactions within liposomal structures.^14^ Cholesterol is included as a crucial component due to its role in enhancing membrane stability and reducing permeability, thus improving drug retention.^14^ Additionally, PEGylated lipids are incorporated to prolong circulation time by providing a steric barrier against opsonization and subsequent clearance by the mononuclear phagocyte system.^14^ Further it assures stability of the liposomes by avoiding the coalescence and fusion of the vesicles.^50^

Liposomes were manufactured following a microfluidic technology as outlined in Fig. 1b. Chips with different channel geometry were 3D printed and utilized in the production offering flexibility and independence at low cost (Supporting information Fig S1).^24^ Immediately following liposome production, the sample underwent analysis via dynamic light scattering (DLS)^52^ and the hydrodynamic diameter and the polydispersity index (PDI) of the liposomes were chosen as the primary response for optimization in this study.^6^

The data set obtained from the lipid screening was analysed using the Tree map (TMAP) dimensionality reduction method (Fig. 2a-c) and an alluvial diagram (Fig. 2d). TMAPs are a tool to inspect high-dimensional datasets in 2-dimensional space while preserving nearest neighbour relationships.^51^ Each point in the TMAP plot corresponds to an instance in the dataset where its location is determined by the underlying formulation parameters such as PEGylated lipids (PEG), cholesterol (Chol), simplified lipid descriptors, and chip geometry (Chip). Further, the colour of each point represents the response value (e.g. Size) of the according data instance. Visualizing this high-dimensional data using TMAP analysis reveals clear clusters corresponding to liposome formation, size, and FRR, indicating that certain combinations of CPPs and lipid formulations consistently produce optimal characteristics. The alluvial plot further illustrates how specific CPPs and lipid formulations influence liposome formation (Fig. 2d and Supporting information Fig S2). This plot shows data flows, representing the number of formed liposomes, between different categories, making it easy to see how variables interact. For example, the cholesterol content appears to affect liposome formation in HSPC and DPPC, as the flows between lipid and Chol %, representing the number of formed liposomes, differ in diameter. However, the plot illustrates the dataset’s complexity, making direct extrapolation impractical and suggesting the use of a ML approach, which can handle large amounts of high-dimensional data and identify patterns not easily seen by traditional methods.

**Fig. 2.**
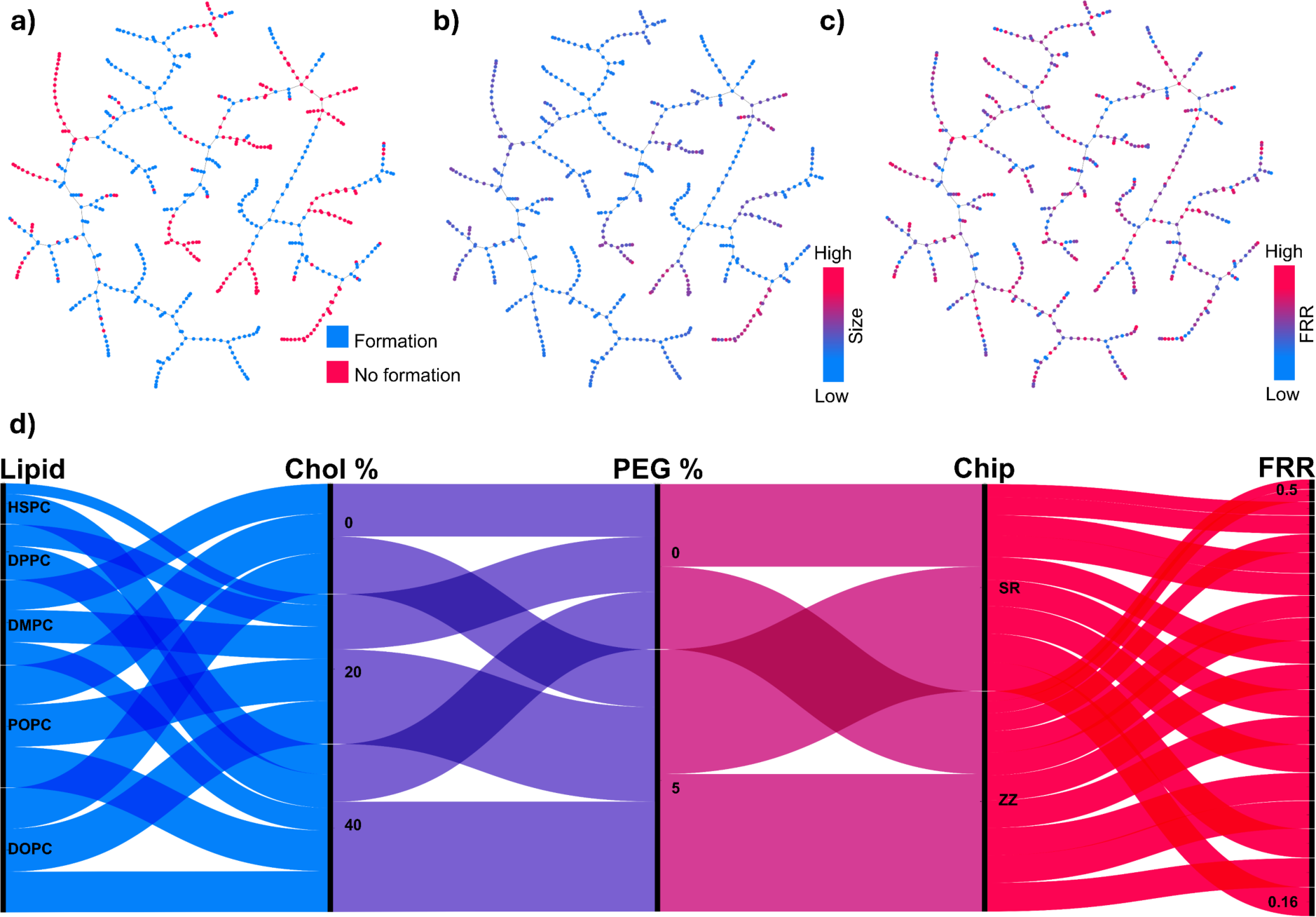
Visualization of liposome data set. a-c) TMAP visualization of the data set for three responses: formation of liposomes (a), size (b), and (c) flow rate ratio (FRR). The features include formulation parameters such as PEGylated lipids (PEG), cholesterol (Chol), and simplified lipid descriptors, and chip geometry (Chip). d) Alluvial plot depicting trends and patterns in the formation of liposomes based on formulation and production parameters. The width of the flows corresponds to the proportion of liposomes in the data set, illustrating the distribution and significance of different parameters.

### Feature selection

A first task in ML is feature selection, which involves identifying and using the most relevant variables (features) in a dataset, associated with microfluidic liposome production. This process reduces overfitting, avoids correlated features that confound the model and improves generalization. The alluvial plot in Fig. 2d supports initial feature selection for further analysis. The selected features include formulation parameters, such as lipid molecular descriptors, cholesterol content, and PEG content, as well as production parameters like chip geometry and FRR. These variables have relevant influences on the process and uniquely contribute to the formation and characteristics of liposomes.^22,24,52,53^ Lipid molecular descriptors provide insights into the physicochemical properties of the lipids, influencing their behaviour in the microfluidic process. While formulation parameters have a direct impact on circulation and drug retention, chip geometry affects the mixing and assembly of lipids, influencing liposome size and uniformity.^24^ The FRR determines the relative velocities of the aqueous and lipid phases, impacting liposome size and encapsulation efficiency.^22,24^ Previous experiments have shown that total flow rate (TFR) becomes less important above a certain high turbulent flow,^27^ where its influence on liposome formation is minimal which is supported by simulations elsewhere.^27,52^ Therefore, the TFR was excluded from the initial feature vector after ensuring to be above the threshold of the used chips (>8 mL/min). A well-constructed feature vector (Fig. 3a) avoids confounding and overfitting, while improving generalization and thereby promises to model the complex underlying data, providing predictive insights and optimizing the microfluidic production process.

**Fig. 3.**
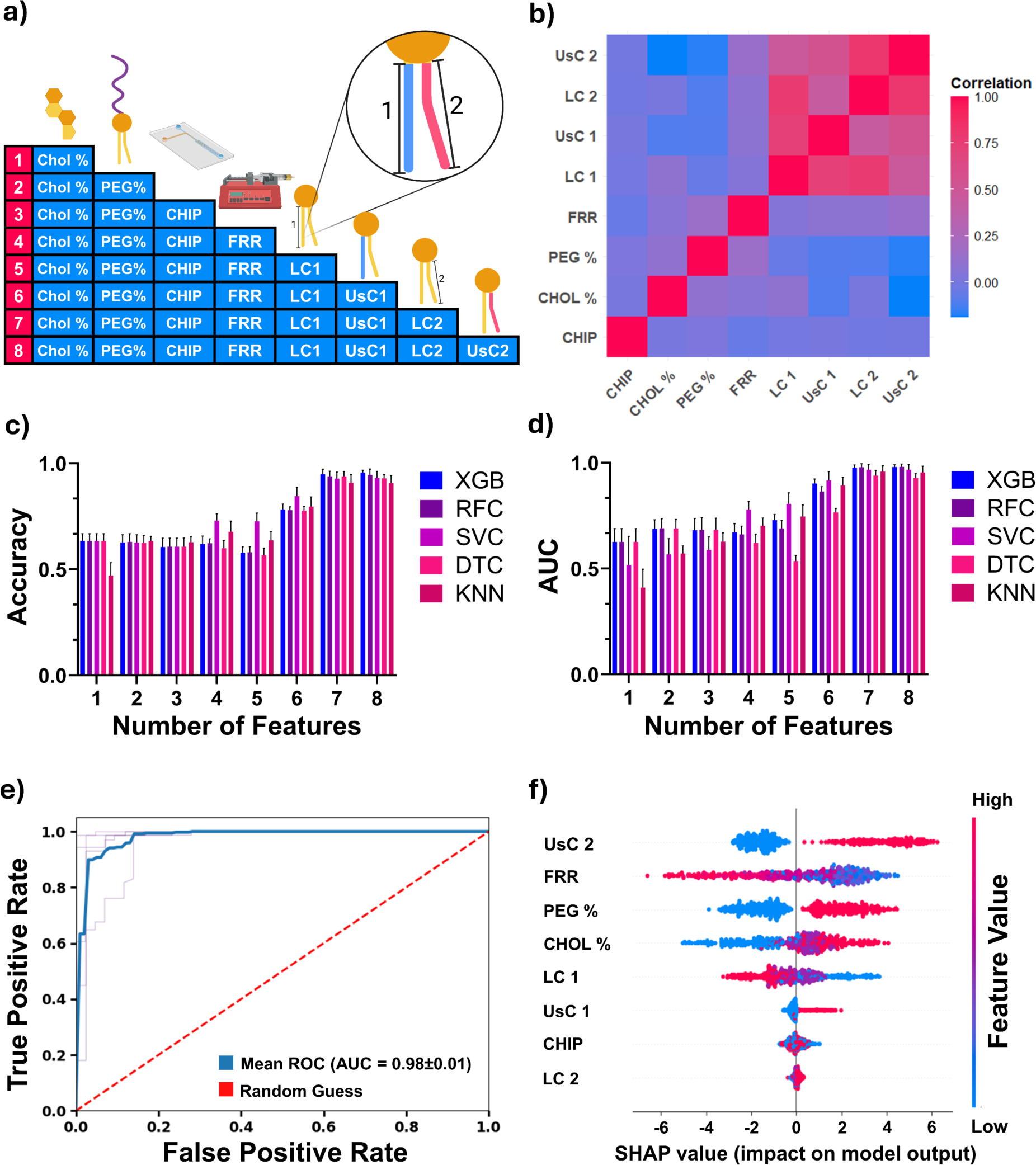
Feature selection and predicting liposome formation: a) Schematic presentation of features used for modelling, including Cholesterol (CHOL), DSPE-PEG percentage (PEG%), Chip geometry, Flow rate ratio (FRR), Length of lipid chains 1 and 2 (LC1/2), and Saturation of lipid chains 1 and 2 (UsC1/2) indicated in blue and magenta respectively. b) Correlation heatmap of all input features. A magenta colour indicates an absolute correlation (= 1) and a blue colour indicates a negative correlation. c) accuracy, and d) Area under the Curve (AUC) of different models for predicting liposome formation, averaged over 5 repeated tests with different data splits (5-fold cross-validation). e) Receiver operator curve (ROC) of 5-fold cross-validation for the XGB model predicting the formation of liposomes. f) SHAP analysis for the 8-feature XGB model illustrating the impact of each feature on the model’s output “liposome formation”, using a swarm plot of SHAP values. The dot colour represents the feature value (magenta for high, blue for low), while horizontal positioning shows the positive or negative contribution of each feature in each prediction instance.

The Spearman correlation matrix provides insights into the correlation between different features in the dataset, independent of the predictive modelling process (Fig. 3b). Each cell in the matrix represents the Spearman correlation coefficient, indicating the strength and direction of the relationship between two features. Positive correlation coefficients indicate a positive linear relationship between features, while negative coefficients signify a negative linear relationship. For example, enhanced correlation coefficients between pairs of features, such as between the lipid descriptors, may indicate multicollinearity, where one feature can be linearly predicted from another which can lead to overfitting. However, in our study, we included these correlated lipid features because, despite their correlation, as literature suggests that multiple lipid characteristics, as captured in our features, are crucial for capturing the underlying microfluidic process.^54^ Additionally, we employed regularization techniques within our models to mitigate any potential issues with multicollinearity, ensuring that the model remains robust and interpretable. Further, tree-based models, less sensitive to multicollinearity, were considered favourable since they allowed us to retain these features without performance loss. Lastly, besides lipid descriptors, the correlations between other features are low, indicating their independence, avoiding overfitting. For example, if two features in the production process model, such as FRR and CHIP, have low correlations it means that changes in CHIP does not affect the FRR. This independence allows the model to accurately understand and predict the effects FRR and Chip have on the outcome without interference. It ensures that each feature contributes uniquely to the model, leading to more reliable and interpretable predictions. In short, a feature vector of 8 process and formulation parameters, relevant to microfluidic liposome production, was chosen for further modelling (Fig. 3a).

## Predicting formation and size of liposomes

### Formation of liposomes

The formation of liposomes during microfluidic production is not always a guaranteed outcome.^55^ It hinges on a complex interplay of formulation and process parameters.^55^ Accurately predicting whether liposomes will form under specific conditions is therefore critical to enriching the pool of successful formulations. To address this challenge, five predictive models based on different underlaying architectures were trained on our dataset generated in-house. The model selection consisted of tree-based models such as decision trees (DT), random forest (RF) and extreme gradient boosting (XGB). Further, models based on support vector classifier (SVC) and k-nearest neighbour (KNN) were trained. Investigating different models allows for a thorough evaluation of their performance and generalization capabilities, facilitating the selection of the most suitable model.^28,29^ In the training process, the feature’s impact on model accuracy, and Area Under the Curve (AUC) of the Receiver Operating Characteristic (ROC) curves were evaluated as seen in Fig. 3c-d. ROC curves are graphical representations that show the performance of a classification model, plotting the true positive rate of a test data set against the false positive rate.^56^ For example, if a model is predicting the formations of liposomes correctly, the false positive rate is low while the true positive rate approaches 1, leading to an AUC of 1, as seen in Fig. 3e. In short, the AUC measures how well a classifier distinguishes between classes, with higher values indicating better performance.^56^

Features were added incrementally (Fig. 3a) in the training process and for each addition, a 5-fold cross-validation was performed. The order of the feature addition was based on reported,^22,24,27,31,46,53^ and experimentally observed influences of certain features during microfluidic production. In short, the feature vector was constructed of formulation features, followed by process features, and finalized with lipid descriptors. Among the models tested, the XGB model, trained on all 8 features, consistently exhibited the highest accuracy, and AUC (Fig. 3e), along with minimal variation between cross-validation sets (Fig. 3c-d). This made XGB the preferred model architecture for the subsequent tasks. Furthermore, tree-based models as XGB are very amenable to model analysis due to their speed which makes XGB an attractive choice for XAI.^29^

The detailed ROC plot for the XGB model, shown in Fig. 3e, illustrates the model’s ability to predict liposome formation with high discrimination power. This is further evidenced by the high AUC scores (Fig. 3e, blue line) consistently achieved by the XGB model across all 5 cross validations. While a functional model is fundamental, it is crucial to learn from the model and derive the findings on the development process.^48^ For example, it can be evaluated what formulation and process features were key contributing factors to the outcome of the model. The importance of input features in predicting liposome formation was determined using Shapley additive explanations (SHAP)^49,57^ analysis for the 8-feature XGB model. In Fig. 3f, the input features are arranged from top to bottom in order of their decreasing impact on the model’s output. The plot highlights the influence of various input features on the model’s predictions and, consequently, potential correlations of these features on liposome formation. For each feature, datapoints are plotted in a dot plot based on their impact on the model prediction (SHAP value) and coloured by their feature value. The plot illustrates the relationship between each feature’s value and its contribution to the model’s prediction, allowing us to understand how changes in the feature value affect the model output and identify patterns such as whether higher or lower values of the feature increase or decrease the prediction. For instance, a high FRR results in a negative contribution to liposome formation, while high PEG content has a positive contribution. Hereby, the SHAP analysis offers insights into potential key drivers influencing liposome formation, such as the degree of unsaturation in the second lipid chain, PEGylation content, and cholesterol content. Conversely, factors like saturation, low FRR, reduced cholesterol content, and lower PEG content might negatively impact liposome formation. It should be noted however, that the SHAP analysis shows the effect of each individual feature on model predictions independently, without considering potential synergies or interactions between input features.

In this section, we validated the XGB model as a robust tool for predicting liposome formation using our in-house dataset. The XGB model exhibited high discriminative power, achieving a mean AUC of 0.98 ± 0.01 across 5-fold cross-validations. Furthermore, the XGB model was suitable for SHAP analysis, an XAI technique that facilitates the analysis of feature importance and their impacts on model predictions. Our analysis highlighted significant influences on the model predictions from factors such as unsaturation of the second lipid chain, PEGylation content, and cholesterol content. These findings suggest that these parameters may play critical roles in liposome formation.

### Predicting size

Having established a reliable model for predicting the formation of liposomes, the next step is to predict their size, a parameter of paramount importance in parenteral drug delivery.^6^ Given the significance of size in determining the therapeutic application, efficacy, and safety of liposomes, developing a model to predict liposome size during microfluidic production is essential. Accurate size prediction facilitates the production of liposomes with tailored properties, enhancing their performance in clinical applications.^6,9,58^

To this end, the previously validated XGB model’s architecture was adapted to a regression model to predict the size of liposomes. By incorporating key formulation and process parameters identified during the formation prediction, the model can be trained to predict liposome size. The selected parameters include lipid structure descriptors, Chol content, PEG content, Chip, and FRR.

To ensure a robust performance assessment, the model was evaluated in a 10-fold cross-validation. In this process, the dataset is split into ten equal parts (folds), with the model iteratively trained on nine folds and tested on the remaining fold. Following cross-validation, the coefficient of determination (R^2^ value) was calculated to quantify the goodness of fit between the predicted and actual particle sizes. An R^2^ value of 0.74 ± 0.08 indicates that approximately 75% of the variance in the particle sizes can be explained by the model. This level of performance can be considered quite reliable in predictive modelling contexts, as it suggests that the model captures a substantial portion of the variability in the target variable. This evaluation framework offers valuable insights into the model’s accuracy, robustness, and generalization ability, supporting informed decision-making in particle size prediction tasks. Further, the relationship between experimentally measured particle size and predicted values is visualized through a scatter plot with a linear regression curve in Fig. 4a. This plot provides insights into the model’s strong predictive capability, with most points clustering around the regression line. Some deviations and outliers, particularly at higher sizes, highlight areas of lesser accuracy. However, these areas are outside the typical size range for therapeutic applications, which is generally between 30-120 nm. In this range, predictive capability was found to be higher (R^2^: 0.78).

**Fig. 4.**
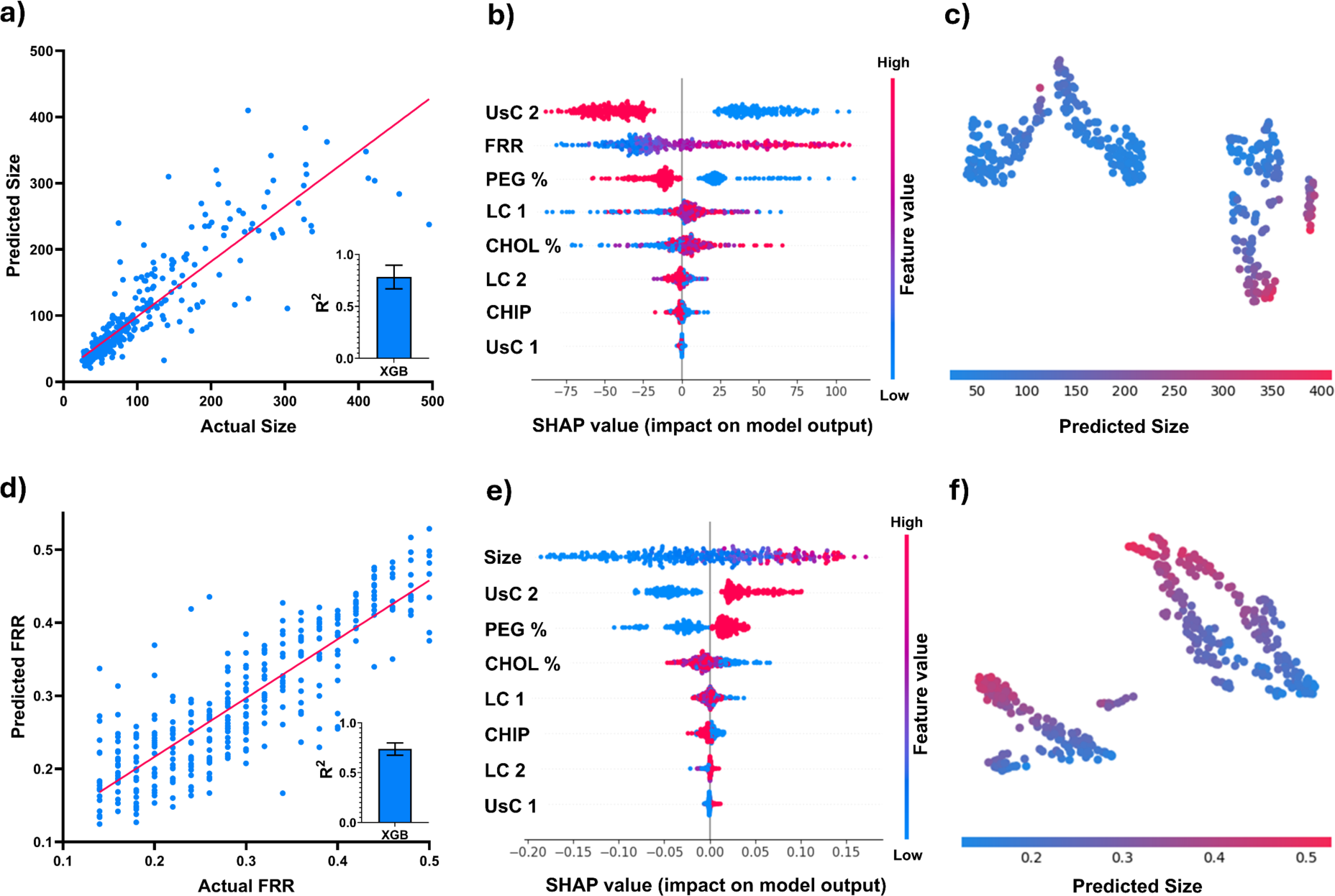
Prediction of the liposomés size and the flow rate ratio (FRR). Model performance depicted as relationship between the experimental a) particle size, or d) FRR, and predicted values. R^2^ value over 10-fold cross-validation quantifies the goodness of fit between the predicted and actual particle sizes or FRR respectively. SHAP analysis for the 8-feature XGB regression model illustrates the impact of each feature on the model’s output b) “liposome size” or e) “FRR”, using a swarm plot of SHAP values. Dot colour represents the feature value (magenta for high, blue for low), while horizontal positioning shows the positive or negative contribution of each feature in each prediction instance. Two-dimensional visualization of the SHAP values calculated for the input features of the XGB model for particle c) size or f) FRR. The SHAP values for the 8 input features were condensed into two principal components using principal component analysis (PCA) and then grouped together using t-distributed Stochastic Neighbor Embedding (t-SNE).

The importance of each input feature for liposome size prediction was investigated via SHAP analysis and visualized in a beeswarm plot (Fig. 4b) which shows the relationship between individual feature values, SHAP values (a measure of feature impact), and predicted sizes. Each point represents an instance in the dataset, with the x-axis displaying the SHAP value for the predicted size. The features are arranged from top to bottom in order of their decreasing impact on the model’s output. This plot reveals how changes in feature values affect predictions, highlighting the importance and influence of different features. Features with a larger spread over the x-axis, such as Saturation of lipid chain 2 (UsC2), Flow rate ratio (FRR), DSPE-PEG percentage (PEG%), Length of lipid chain 1 (LC1), and Cholesterol content (CHOL%), may have a more significant impact on the model’s predictions. These are potential key formulation and production factors that determine the size of a liposome. For instance, a high (magenta) degree of unsaturation in the lipid chain 2 and a high PEG content seem to have a negative contribution to the size output while high FRR, the length of lipid chain 1 and the cholesterol content do the contrary. Further clusters or patterns, as for UsC2 and PEG%, in the plot reveal groups of instances with similar feature contributions and predicted sizes. This implies that the model identifies lipids with unsaturated chains and formulations with high PEG content along with FRR as significant influences on the liposomés size prediction. Overall, the plot illustrates that while certain formulation and process parameters significantly impact size prediction, the process is undeniably a complex interplay of various factors. Consequently, it is evident that liposome size depends on the interaction of all these parameters, as supported by literature.^16,22,23^

For enhanced interpretability, high-dimensional SHAP values are reduced using Principal Component Analysis (PCA) as shown in Fig. 4c. The SHAP values for the 8 input features are condensed into two principal components and visualized in a scatter plot. Each point represents an instance in the dataset, positioned by the t-distributed Stochastic Neighbour Embedding (t-SNE) algorithm, and coloured according to the predicted size. Clusters of nearby points indicate instances with similar feature contributions, revealing patterns within the data. Colour coding highlights regions with similar predicted sizes, providing further insights into prediction patterns. Isolated or distant points may indicate anomalies or unique feature contributions. Larger sizes (magenta) are clearly clustered, indicating that the model effectively distinguishes between particle sizes.

Overall, we demonstrate a robust model predicting liposome size with high predictive capability (R^2^: 0.78). SHAP analysis yields practical insights, such as identifying influential formulation and process features that clarify the model’s decision-making process and key factors driving predicted sizes. Evaluating prediction variations across segments validates the model’s performance and its ability to capture underlying data patterns.

### Models to predict microfluidic process parameters

Translating liposome production from the laboratory to clinical and industrial applications poses significant challenges.^17,59,60^ Scaling up production while maintaining consistency and reproducibility is a primary concern.^17^ With traditional liposome production methods often failing to scale effectively, prompting a shift towards microfluidic techniques due to their sustainability and applicability in industry.^15^ However, this transition is not straightforward. Each liposomal formulation can respond differently to microfluidic processes, necessitating extensive trial and error and laborious optimization on a case-to-case basis.^15^ Furthermore, variations in microfluidic chip design complicate the standardization of production processes.^52^ Robustly 3D-printing these chips might offer a pathway to standardization, facilitating broader application in pharmaceutical manufacturing.^24^ Additionally, microfluidic production depends on many controllable parameters (Formulation, Chip, FRR, etc.), but this makes the design space high-dimensional and impractical to empirical testing. ML can help to efficiently navigate the high-dimensional design space and help us find the correct parameters to produce liposomes with wanted properties in a microfluidics production. A predictive model that takes formulation parameters and chip geometry into account to determine the optimal FRR for microfluidic production, targeting desired CQA like size, can significantly ease the transition to microfluidics and scale-up. By accurately predicting FRR, such a model can streamline development, reducing the need for extensive empirical optimization. Building on previous analyses, key features influencing liposome formation and size have been identified. Namely, formulation parameters such as unsaturation and length of the lipid chains, cholesterol and PEGylation content along with the liposome’s desired size and the used chip geometry. These features serve as the foundation for developing an XGB model to predict the most critical process parameter, FRR, ensuring comprehensive optimization of the microfluidic process.

The relationship between the experimental and predicted FRR values for a given liposome batch is visualized in a scatter plot (Fig. 4d), showing a linear trend, implying that the predicted FRR for a certain liposome batch is in accordance with the experimental. The model’s performance robustness is further evaluated using 10-fold cross-validation, involving 10 repeated tests with different data splits. Following cross-validation, the R^2^ value of 0.74 ± 0.06 indicates that the model explains approximately 75% of the variance in FRR, demonstrating reliable performance in predicting the FRR. A beeswarm plot (Fig. 4e) of calculated SHAP values reveals the impact of the feature “liposome size” as the main driver for FRR prediction before the unsaturation of lipid chain 2 (UsC2). It is evident that larger liposomes, high UsC2, and PEG% along with a low Chol % have a positive contribution to the model’s output. This might imply that both, chemical formulation and morphological factors play a major role in selecting a FRR to obtain a desired liposome in a microfluidic process. Further, high-dimensional SHAP values are reduced using PCA and t-SNE (Fig. 4f) and visualized to reveal data patterns and segments. It is visible that clear gradients of FRR ratio span through the clusters, implying the model’s ability to making decision and thereby further justifies the feature vector.

This robust predictive model for FRR has the potential to ease the transition to microfluidic production of liposomes, ensuring consistent and scalable manufacturing processes. By accurately predicting FRR, this model reduces dependency on empirical optimization, accelerating the transition from bench-scale experiments to industrial production. This advancement holds immense potential to streamline the development of liposomal drug delivery systems, facilitating their journey from the laboratory to clinical and industrial settings. It represents a significant step forward in optimizing these systems, promising improved therapeutic outcomes and broader accessibility.

### Wet lab validation

The last step involved validating our XGB models in real-world conditions, and assessing how effectively these models transition from theoretical development to actual liposome production. To validate the models, a transition workflow from traditional to microfluidic liposome production was simulated, employing predictive models validated through a series of experiments as illustrated in Fig. 5a. The validation process involved randomly selecting three distinct liposome formulations, each assigned a specific desired size. Additionally, a specific chip geometry was chosen, engineered, and printed in an independent laboratory.

**Fig. 5.**
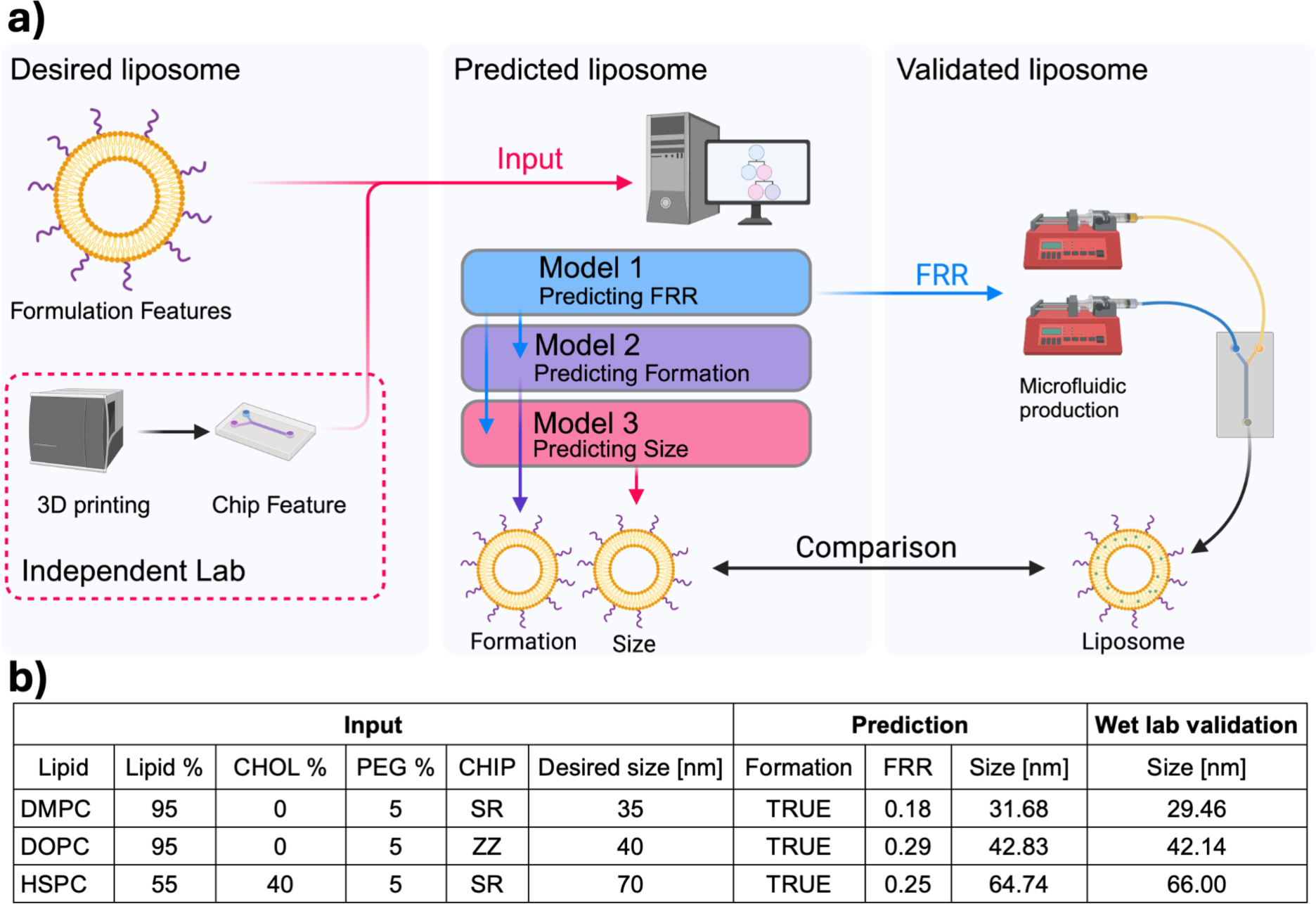
Transition workflow for validation experiments. a) desired liposome and lab set up was fed to model to predict process parameters which were fed to consecutive models to predict CQAs such as the formation and size of liposomes which was compared to liposomes from microfluidic production. b) Wet lab validation for 3 liposomes with desired input formulation and size, predicted CQAs and FRR and according liposomes produced in the wet lab validation.

The workflow starts with a model predicting the FRR necessary for each formulation. Subsequently, a second model assessed whether the liposomes would form under the specified process conditions for the given lipid formulation. Finally, a third model predicted the resulting liposome size. After confirming the predictive accuracy of the model, the liposomes were produced using the predicted process parameters. The sizes of the produced liposomes were then measured via DLS and compared to the predicted and desired sizes. The results in Fig. 5b demonstrated a deviation of approximately 4 nm, indicating that the model’s predictive power is favourable. This is further supported by the performance relationship plots, illustrating the performance along the transition workflow (Supporting Information Fig. S3a-c).

This validation process highlights the model’s robustness in predicting critical parameters for liposome formation and size, ensuring a seamless transition from traditional to microfluidic production methods. Moreover, the model demonstrates exceptional value in adapting existing liposomal formulations to microfluidic processes, significantly streamlining the development of advanced liposomal drug delivery systems.

## Conclusion

In this study, we demonstrated that the integration of predictive modelling with wet lab validation underscores the potential of ML for precise and efficient optimization of microfluidic liposome production, capable of enhancing the development effort, scalability and transitioning to microfluidic technology in pharmaceutical sciences. This approach not only streamlines the production process but also paves the way for similar models in drug encapsulation within liposomes and the expansion to commercial microfluidic chips. Future research could explore the application of such models in also optimizing drug release profiles, ensuring that liposomal formulations deliver therapeutic agents effectively and consistently. The advancements in ML-driven microfluidic production hold promise to accelerate the development of drug delivery systems, making them more accessible and adaptable to a wide range of pharmaceutical applications.

## Materials and methods Materials

The phospholipids DMPC (1,2-dimyristoyl-*sn*-glycero-3-phosphatidylcholine), DOPC (1,2-dioleoylglycerol-3-phosphorylcholine), DPPC (1,2-dipalmitoyl-*sn*-glycero-3-phosphatidylcholine), HSPC (hydrogenated soy phosphatidylcholine), DSPE-PEG2000 (1,2-distearoyl-*sn*-glycero-3-phosphoethanolamine-N-[methoxy(polyethylene glycol) monosodium salt), POPC (1-palmitoyl-2-oleoyl-glycero-3-phosphocholine) were purchased from Lipoid, Germany. Cholesterol was purchased from Sigma Aldrich, USA. Purified water was obtained using a Barnstead Smart2Pure system from Thermo Fisher Scientific Inc., Germany. PBS (phosphate buffered saline, pH 7.4) was obtained using KCl (potassium chloride), KH_2_PO_4_ (Potassium Dihydrogen Phosphate), NaCl (sodium chloride), NaH_2_PO_4_ (Sodium Phosphate Monobasic), Na_2_HPO_4_ (Sodium Phosphate Dibasic), NaOH (sodium hydroxide) from Carl Roth LLC, Germany. NaOH (sodium hydroxide in pellet form) from Haenseler, Switzerland. Absolute Ethanol (EtOH) was purchased from VWR International SAS, France. Ethanol denatured with ketone (EtOH 94%) was purchased from Dr. Grogg Chemie AG, Switzerland. Methanol (MeOH) was purchased from Fisher Chemical, Belgium.

## Methods

### Data collection

For the liposome screening, formulations were collected following a Design of Experiment ensuring evenly distributed data points. A full factorial design plan was created on Minitab 2018 (Minitab Inc., USA) for each considered lipid including the factors, Chip, FRR, TFR, cholesterol and PEG content. For the screening, lipid masses were calculated, weighed, and dissolved in ethanol (≥99.8% v/v) at a total lipid concentration of 5 mg/mL. The aqueous buffer used was PBS with a pH of 7.4. For the preparation of liposomes via microfluidics, the chip, positioned in a 55 °C water bath, was connected to two syringes, which were mounted on two NE-1010 Higher Pressure Syringe Pump, KF TECHNOLOGY SRL, Italy. The connection was established through polyethylene tubing. A specific amount of lipid was delivered against PBS at controlled flow rates. Various amounts of DSPE-PEG 2000, ranging from 0% to 5%, and Cholesterol, ranging from 0%-40%, were added to the lipids and were included in the screening. The two syringes containing the respective phases were preheated to a temperature of 62 °C to remain above the phase transition temperature. The microfluidic chips from the University of Urbino, Italy, were manufactured using ultrafuse polypropylene (PP) with an Ultimaker 3D printer from Ultimaker in The Netherlands (See Supporting information, Fig S1). After discarding the initial 1 mL from the outlet of the chip, the resulting samples were collected.

The chip cleaning process involved sequential flushing steps, each carried out consecutively (10 mL of NaOH 1 M, 10 mL of NaOH 0.1 M, 20 mL of MQ Water, 10 mL of EtOH 94%). Upon the conclusion of the procedure, the chip was dried and flushed using 6 mL of methanol. 1 mL of methanol was collected and analysed via HPLC CAD^62^ to validate the process and confirm the chip’s cleanliness with UltiMate 3000 HPLC from Thermo Fisher Scientific Inc., Germany (Supporting Information, Fig S4). To conclude the cleaning process, the chip was washed with 10 mL of EtOH 94% and dried.

Immediately following liposome production, the sample underwent analysis via Dynamic Light Scattering (DLS). DLS was utilized to evaluate the population, size, and polydispersity index (PDI) of liposomes using the Litesizer DLS 500 (Anton Paar GmbH, Austria) with a 173° backscatter angle and a 633 nm helium–neon-laser. Particle size measurements were performed in PBS (Refractive index: 1.335) at a pH of 7.4 and a temperature of 25°C. The measurements were corrected to the viscosity of the according sample. After equilibrating the sample, the measurement (10 runs × 10 s) was performed. If the intensity size distribution of the liposome was unimodal, the autocorrelation function was analyzed according to the cumulant method by the Kalliope™-software. For our model we considered samples below 500 nm and a monodisperse size distribution as forming liposomes.

### Data curation

The dataset used for ML model development underwent a preprocessing step to ensure data quality and consistency. Initially, any observations where the ‘Size’ variable exceeded 500 nm were filtered out to focus the analysis on liposomes within a specific size range. Also, samples with PDIs above 0.3 were considered as “no formation” as the consensus in parenteral drug delivery. The remaining data was then prepared for model training and evaluation. Additionally, observations associated with a multidisperse size distribution (Population>1) were excluded from the dataset for size and FRR modelling.

### ML Model Development, Evaluation and Interpretation

The ML models were given the input features comprising of ‘CHOL %’, ‘DSPE-PEG %’, ‘FRR’, and parameters related to chain length and unsaturation. The predictive process initiates with the transformation and standardization of feature data, ensuring its readiness for modelling. Categorical variables, such as the ‘CHIP’ parameter, were one-hot encoded to convert them into a suitable format for model training. For the models trained to predict the FRR the feature ‘FRR’ was replaced by ‘Size’. Python module sckit-learn and xgboost were mainly used for modelling.

The performance of the developed ML model was assessed using a stratified K-fold cross-validation approach from the Scikit-learn library in Python for the classification models and a K-fold cross validation for the regression models (10:90; Test:Train). For classification models accuracy, precision and AUC were determined. Regression models were evaluated by assessing their R^2^ value. Additionally, SHAP^49^ (SHapley Additive exPlanations) analysis was conducted to interpret the model’s predictions and understand the relative importance of each input feature. SHAP values were computed to quantify the contribution of each feature to the predicted output. Visualization techniques, including SHAP summary plots and scatter plots, were employed to visualize the relationships between predicted and actual liposome sizes, providing insights into the model’s performance and feature importance. Furthermore, Spearman’s correlation heatmap were generated to explore the correlation between input features and liposome size. Lastly, unsupervised clustering techniques, including PCA (Principal Component Analysis) and t-SNE (t-distributed Stochastic Neighbor Embedding), were utilized.

### Validation

The validation process involved randomly selecting three distinct liposome formulations, each assigned a specific desired size. Additionally, a specific chip geometry was assigned and printed. Microfluidic chips were printed in an independent lab using the 3D printer Stratasys Eden260VS (Stratasys, Germany) and the photopolymer IORA Model White (iSquared2, Switzerland). Predictive models were tasked to predict the FRR, formation and the resulting liposome size. The liposomes were produced using the predicted process parameters. The size of the produced liposomes was then measured via DLS and compared to the predicted and desired sizes.

## Statistics and reproducibility

All statistical analyses performed are available for each experiment in the results & discussion section of the manuscript. No data were excluded from the analysis. GraphPad Prism 10.2.3 (GraphPad Software, USA) and R 4.4.0 (The R Foundation, Germany) were used for statistical analysis.

## Data availability

All data generated or analyzed during this study are included in this published article and its supplementary information file or are available from the corresponding authors upon reasonable request. Source data are provided with this paper.

## Code availability

The codes that support the findings of this study are available on GitHub (https://github.com/Luciani-Group/micro_fluidic_Liposome). Code was written and executed in Python 3.11.9 in an Anaconda 24.3.0 environment.

## Author contributions

RE: Conceptualization, Methodology, Investigation, Validation, Visualization, Formal analysis, Writing – original manuscript.

MO: Methodology, Investigation, Visualization, Formal analysis, Review & editing

GB: Methodology, Investigation, Validation, Review & editing

NS: Investigation, Validation

MT: Methodology, Review & editing

LC: Methodology, Funding acquisition, Review & editing

JLR: Methodology, Funding acquisition, Review & editing

SA: Conceptualization, Methodology, Supervision, Data curation, Review & editing

PL: Conceptualization, Supervision, Project administration, Funding acquisition, Review & editing.

## Acknowledgement

Prof. Dr. Manuela Eugster from the ARTORG Center for Biomedical Engineering Research at the University of Bern, for printing the chips used for the wet lab validation.

Fig. 1 & 5 was created with Biorender.com.

This work has been partially funded by the European Union – NextGenerationEU - under the Italian Ministry of University and Research (MUR) National Innovation Ecosystem grant ECS00000041 – VITALITY - CUP [H33C22000430006].

