## Supporting Information for "Leveraging machine learning to streamline the development of liposomal drug delivery systems"

**Affiliations:**

### Supporting information

#### Microfluidic chip

The 3D printed chips used in this work were previously developed at the University of Urbino Carlo Bo.[1] Briefly, the chips present two different designs that allow an effective micromixing of the fluids in the channels of both. The ZZ design presents a zigzag bas-relief (Fig. S1A) meanwhile the SR design present a split and recombine asymmetrical and circular path (Fig. S1B). For both the chips, the inlets have a 1mm squared shape that become 0.4mm and 0.6mm wide in the asymmetric channels of the SR chip meanwhile in the ZZ chip, the bas-relief have a high of 0.5mm. The microfluidic devices were produced using a Fused Deposition Modeling 3D printer (Ultimaker 3, Ultimaker, The Netherlands) with a polypropylene filament to ensure the higher solvent and chemical compatibility.

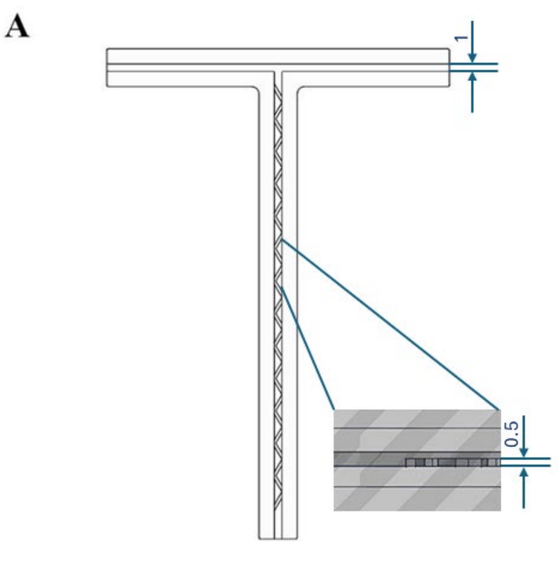


**Fig S1** Computer aided design models of A) “ZZ” (zigzag design) microfluidic chip and B) “SR” (split and recombine design) microfluidic chip.

**Data set**

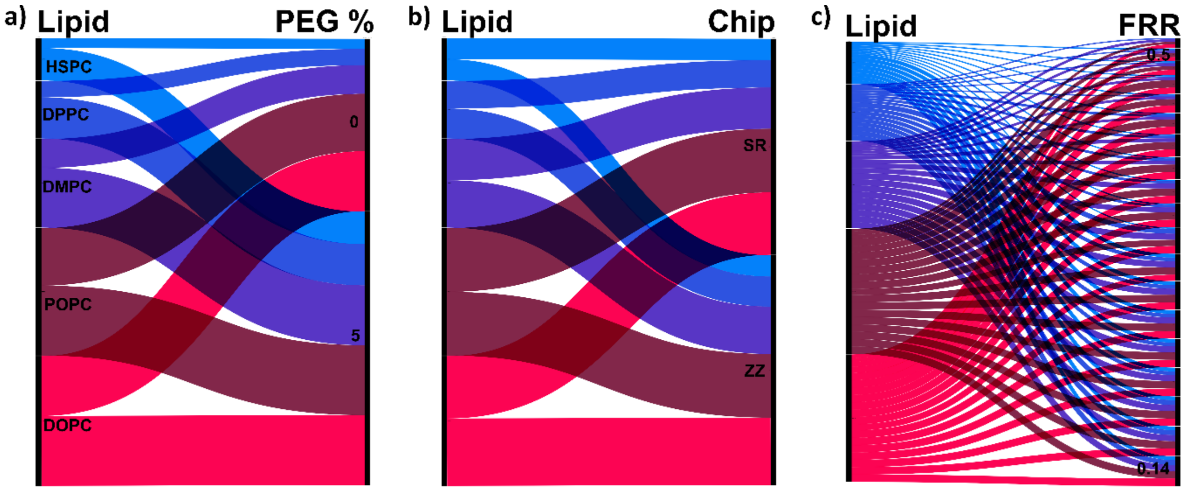


Fig S2 Alluvial plot depicting lipid trends in forming liposomes. a) Formation of Liposomes originating from lipids to PEG % content. B) Formation of Liposomes originating from lipids to Chip geometry. C) Formation of Liposomes originating from lipids to FRR.

### Validation of transitioning workflow

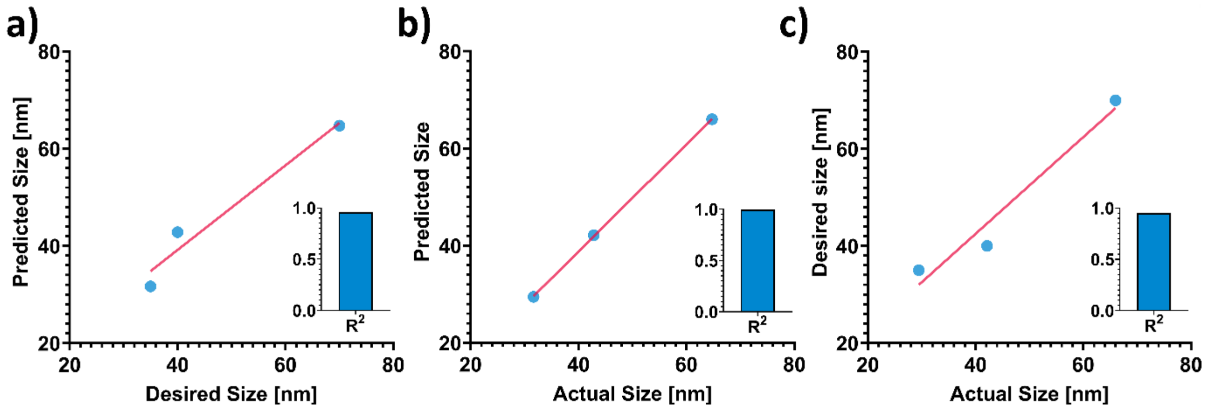


Fig S3 Performance relationship for transitioning work flow b) Model performance depicting desired size vs. predicted size after CPP feedback. c) and actual size of liposomes after production vs. predicted size. d) Front to end performance visualizing desired size vs. actual size.

### Microfluidic chip cleaning

The chip cleaning process involved sequential flushing steps, each carried out consecutively (10 mL of NaOH 1 M, 10 mL of NaOH 0.1 M, 20 mL of MQ Water, 10 mL of EtOH 94%). Upon the conclusion of the procedure, the chip was dried and flushed using 6 mL of methanol. 1 mL of methanol was collected and analysed via HPLC CAD[2] to validate the process and confirm the chip's cleanliness with UltiMate 3000 HPLC from Thermo Fisher Scientific Inc., Germany (Supporting Information). To conclude the cleaning process, the chip was washed with 10 mL of EtOH 94% and dried. HPLC analysis of lipid standards and samples collected before chip usage and after washing, seen in Fig S2, validate the cleaning procedure.

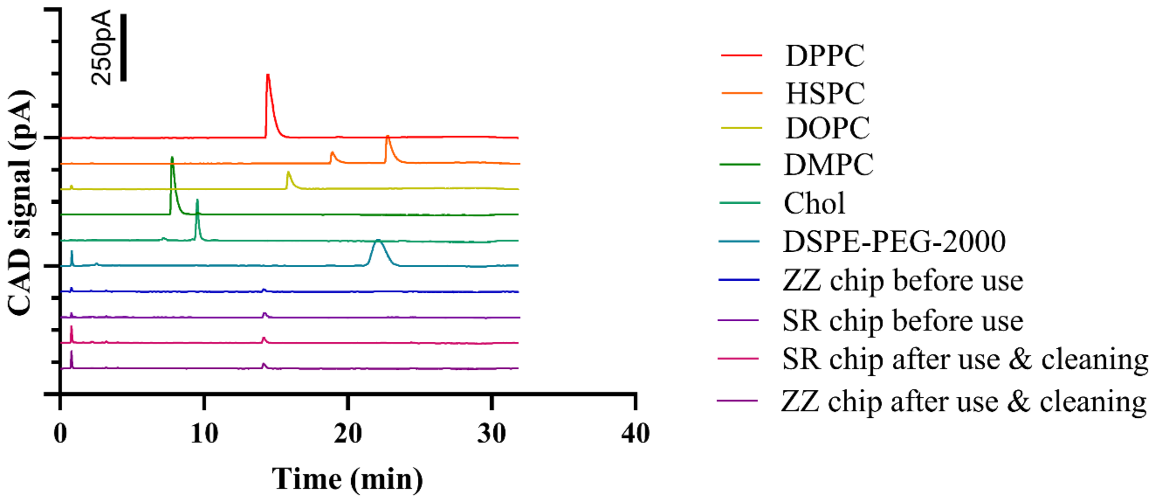


Fig S4 HPLC Cad analysis of lipid standards and methanol samples collected from the chips before use and after use and cleaning procedures.
